# Uncovering the structural flexibility of SARS-CoV-2 glycoprotein spike variants

**DOI:** 10.1101/2022.04.20.488873

**Authors:** Hiam R. S. Arruda, Tulio M. Lima, Renata G. F. Alvim, Fernanda B. A. Victorio, Daniel P. B. Abreu, Federico F. Marsili, Karen D. Cruz, Patricia Sosa-Acosta, Mauricio Quinones-Vega, Jéssica de S. Guedes, Fábio C. S. Nogueira, Jerson L. Silva, Leda R. Castilho, Guilherme A. P. de Oliveira

## Abstract

The severe acute respiratory syndrome CoV-2 rapidly spread worldwide, causing a pandemic. After a period of evolutionary stasis, a set of SARS-CoV-2 mutations has arisen in the spike, the leading glycoprotein at the viral envelope and the primary antigenic candidate for vaccines against the 2019 CoV disease (COVID-19). Here, we present comparative biochemical data of the glycosylated full-length ancestral and D614G spike together with three other highly transmissible strains classified by the World Health Organization as variants of concern (VOC): beta, gamma, and delta. By showing that only D614G early variant has less hydrophobic surface exposure and trimer persistence at mid-temperatures, we place D614G with features that support a model of temporary fitness advantage for virus spillover worldwide. Further, during the SARS-CoV-2 adaptation, the spike accumulates alterations leading to less structural rigidity. The decreased trimer stability observed for the ancestral and the gamma strain and the presence of D614G uncoupled conformations mean higher ACE-2 affinities when compared to the beta and delta strains. Mapping the energetic landscape and flexibility of spike variants is necessary to improve vaccine development.

## Introduction

Pathogenic coronaviruses (CoVs) have crossed species to cause pneumonia. CoV causing severe acute respiratory syndrome (SARS-CoV) emerged in 2003 in Guangdong Province, China, and caused more than 700 deaths in 5 countries. The Middle-East respiratory syndrome CoV (MERS-CoV) appeared in 2012 in the Arabian Peninsula, causing 858 known deaths over the last ten years, at a high case fatality rate, estimated at 35% of patients reported with MERS(1, 2). On the other hand, the severe acute respiratory syndrome CoV-2 (SARS-CoV-2), first identified in December 2019 in the Wuhan Province, China, rapidly spread worldwide. The sharp escalation forced the World Health Organization (WHO) to declare a pandemic in March 2020. Within two years of its identification in Wuhan (as of Dec 2021), it had caused more than 270 million cases with 5.3 million deaths worldwide. Four other endemic CoVs of low pathogenicity affect humans: hCoV-OC43, HCoV-HKU1, HCoV-NL63, and HCoV-229E. They are seasonal viruses accounting for ∼15% of common colds(3).

The sequence of the SARS-CoV-2 harbors 96% similarity with the CoV RaTG13 of bats and lower percentages compared to SARS-CoV (79.5%) and MERS-CoV (55%). It indicates that bats may have been the reservoir hosts for the emergence of SARS-CoV-2(4, 5). The idea of SARS-CoV-2 as a “generalist virus” is based on positive selection acting at the base of the bat lineage that it emerged from(6). Direct transmission from bats to humans or the presence of an intermediate host is a matter of discussion still(1). SARS-CoV-2 has peculiarities from other β-CoVs: *(i)* a polybasic furin protease-like (RRAR/S) site between the S1 and S2 subunits of the spike glycoprotein(7); *(ii)* the presence of novel O-glycan binding sites flanking the protease cleavage site in the spike(8).

The 2019 CoV disease (COVID-19) has a broad clinical and epidemiological profile(9). Some have mild to moderate symptoms, including fever, cough, and breathing difficulty, and many are asymptomatic. However, a relatively high percentage of infected individuals develop severe manifestations, including respiratory failure requiring mechanical ventilation and aggressive inflammatory response. Debilitating symptoms are fatigue and dyspnea seen up to two months after the viral load decreases(10). Individuals over 50 years old and with chronic conditions have an increased probability of evolving severely with a risk of death.

The spike glycoprotein is the leading glycoprotein at the viral envelope and the primary antigenic candidate for vaccines against COVID-19. Each protomer has two subunits, S1 (residues 14-685) and S2 (residues 686-1273), in addition to a signal peptide (residues 1-13). Essential domains of S1 are the N-terminal domain (NTD, residues 14-305) and the receptor-binding domain (RBD, residues 319-541) (Figure 1a). S2 carries the fusion peptide (FP, residues 788-806), repetitive heptapeptide sequences (HR1, residues 912-984; HR2, residues 1163-1213), the transmembrane domain (TM, residues 1213-1237), and the connector domain (CD, residues 1237-1273) (Figure 1a). The RBD binds to the host receptor, the peptidase angiotensin-2 converting enzyme (ACE 2)(7). Cryo-EM revealed that H-bonds and salt bridges govern RBD-ACE 2 interactions(11). After the interaction, the S2 subunit undergoes conformational changes exposing the FP to interact with the host membrane. It leads to HR1-HR2 interactions bringing the viral envelope close to the cell’s surface(2), and fusion occurs when host proteases promote the cleavage of the furin-like site. Weak hydrophobic interactions drive the trimer interface, unlocked during pre/postfusion conformational positioning(12).

**Figure 1.**
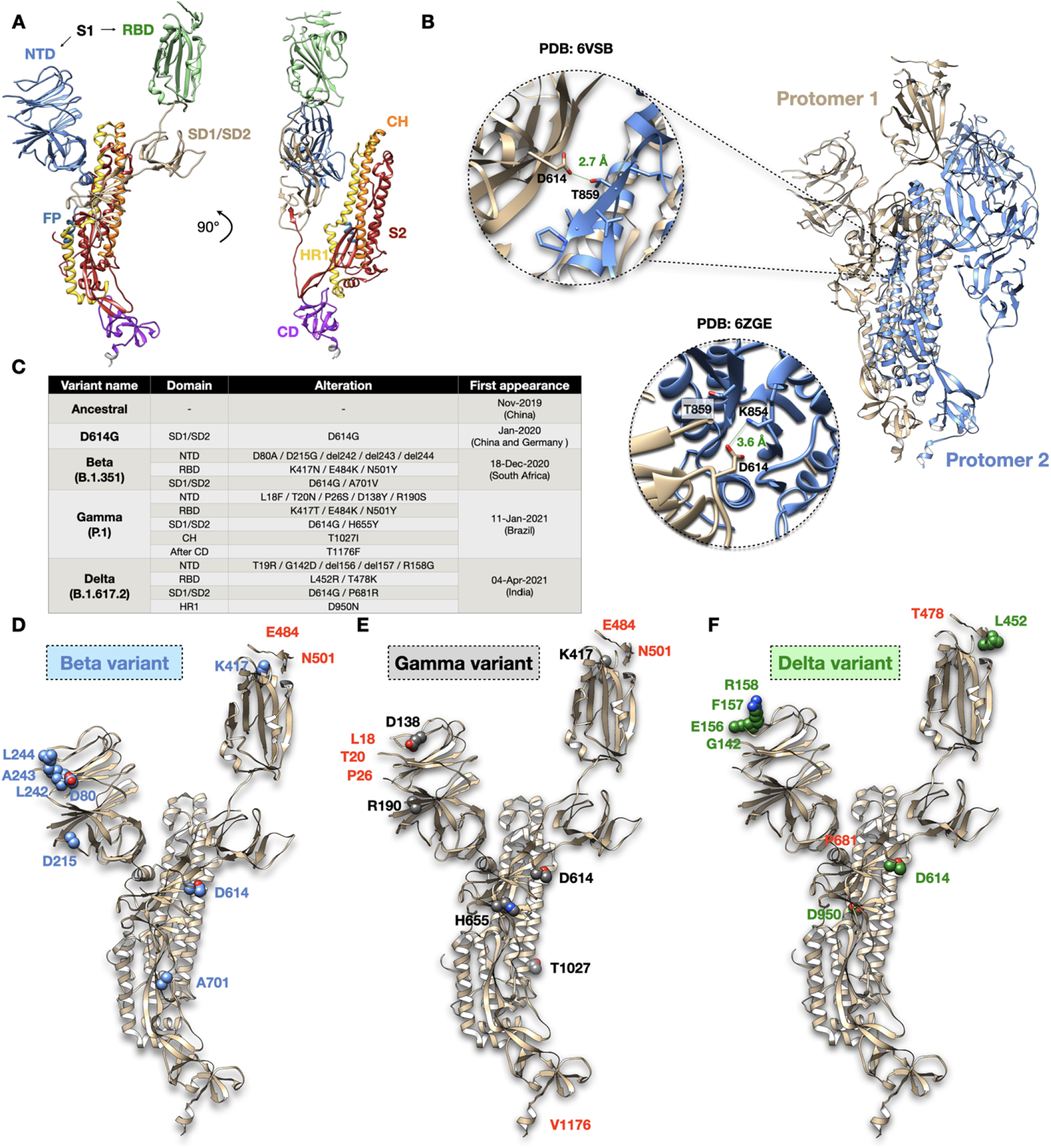
Spike glycoprotein and the variants of concern. **A**, Atomic model (PDB code: 6VSB) showing the ribbon representation of the spike protomer obtained from cryo-EM maps. Schematics is color-coded for the N-terminal domain (NTD, blue); receptor-binding domain (RBD, green); sub-domain 1 and 2 (SD1/SD2, tan); fusion peptide (FP, turquoise); central helix (CH, orange); heptad repeat (HR1, gold yellow); S2 subunit (dark red); connector domain (CD, purple). The NTD and RBD form the S1 subunit. **B**, Atomic model highlighting two spike protomers and the D614 interprotomer interaction. Zoomed images show D614-T859 H-bond interaction (PDB code: 6VSB) at 2.7 Å distance and the D614-K854 salt bridge at 3.6 Å distance (PDB code: 6ZGE). **C**, The table shows features of studied spikes (ancestral strain, D614G, beta, gamma, and delta variants). The altered residues of each variant, their domain locations, and the date and place of the first appearance are shown. **D-F**, Atomic model (PDB code: 6VSB) showing altered sites for **(D)** beta variant in blue, **(E)** gamma variant in black, **(F)** delta variant in green. Altered sites are shown as spheres. Labels in red refer to residues not resolved on the cryo-EM map.

Virus evolution is an issue of concern during a pandemic. Although most viral mutations are expected to be either harmful, swiftly purged, or neutral, a small proportion may alter the virus biology, affecting pathogenicity, infectivity, and antigenicity(13). After a period of evolutionary stasis, a set of SARS-CoV-2 mutations has arisen as “variants of concern” (VOC)(6). The initial months of the COVID-19 outbreak featured a randomly neutral and weakly purifying selection as expected at the early phases of transmission(6). At this stage, a naive susceptible population is not exerting any selective pressure on the pathogen(6). Fitness-enhancing mutations such as the D614G in the spike(14, 15) and the P323L in the RNA-dependent RNA polymerase (RdRP)(16) emerged, contributing to viral transmissibility but fewer to pathogenicity(17). Other early detected mutations include N439K and Y453F in the spike, leading to enhanced RBM-ACE 2 affinity(18). N439K circulated widely in European countries (lineage B.1.258), while Y453F has been identified in Denmark over sequences related to infections in humans and minks (lineage B.1.1.298)(19).

Wide-genome search aiming to find signatures for adaptive pressure captured an increase in selective forces acting on SARS-CoV-2 genes(20). Adaptive pressure is a natural consequence presumably because of increasing host immunity due to infection or vaccination and the shift in the host environment due to public health intervention and social distancing. This positive selection changed the virion antigenic phenotype, and variants emerged carrying more convergent mutations that compromised the host immune recognition. In this vein, three divergent SARS-CoV-2 lineages succeeded, namely the alpha (B.1.1.7), the beta (B.1.351), and the gamma (P.1) variants(21, 22).

Known as 501Y lineages due to the persistence of an N501Y substitution in the RBD, these variants presented altered phenotypes, including increased ACE-2 affinity(23), transmissibility, and capacity to overcome vaccination-induced immunity(24). E484 is an immunodominant site with various substitutions, including the E484K present within the beta and the gamma. Together, N501Y and E484K increased RBD-ACE-2 affinity to 12.7 fold(23) and decreased neutralization by convalescent sera (25), vaccine-elicited antibodies, and monoclonal antibodies(26, 27).

Several mechanisms have been considered to affect antigenicity. The most common one is footprint amino acid substitutions altering the epitopes and consequently the neutralizing activity. Other examples include increasing the receptor-binding avidity, deletions and insertions, transforming glycosylation sites, and the presence of allosteric effects(17). Within the context of glycosylation changes, the T20N substitution in the gamma variant introduced a new glycosite(28). The lack of extensive glycosylation in RBD and its abundance in NTD has been hypothesized to play a role in RBD immunogenicity(29, 30). In this vein, 41 monoclonal antibodies (mAbs) recognized the NTD and neutralized SARS-CoV-2, allowing the definition of an antigenic NTD map(30). The presence of several substitutions, additions, and deletions within the NTD indicates the domain is under selective pressure and should compose the targetable toolkit to avert SARS-CoV-2 progression.

While undergoing antigenic evolution, one fundamental question is whether authorized vaccines will maintain protection from severe disease and death for future SARS-CoV-2 variants. The temporal and geographic efficacy is an aspect requiring constant surveillance. Updated vaccines tailored to emerging antigenic variants that cross-react against several circulating variants are one of the possible approaches to fight the pandemic. Here, we present comparative biochemical and biophysical data of the full-length ancestral and D614G spike together with three other highly transmissible strains that the WHO classified as variants of concern (VOC): beta, gamma, and delta. We found evidence to argue that during the SARS-CoV-2 adaptation, the spike glycoprotein accumulates alterations that lead to a tendency of less structural rigidity.

Further, we revealed D614G features supporting a model of temporary fitness advantage required for virus spillover worldwide but not for long-term adaptation. The decreased trimer stability observed for the ancestral and the gamma strain and the presence of D614G uncoupled conformations mean higher ACE-2 affinities when compared to the beta and delta strains. Mapping spike variants’ energetic landscape and flexibility are necessary to improve vaccine development.

## Results

### The glycosylation pattern of SARS-CoV-2 spike variants

We produced the full-length spike containing the ancestral sequence, the D614G early variant, and three highly transmissible variants that the WHO classified as variants of concern (VOCs): beta, gamma, and delta (Figure 1b-f). We performed quality checks to ensure our preparations’ identity and purity (Figure 2a and Supplementary figures 1 and 2). A sandwich ELISA confirmed that the proteins were functional and bound to the ACE-2 receptor, and to a monoclonal antibody developed against the ancestral spike protein (Supplementary figure 3). We confirmed a peptide coverage higher than 94% of studied variants (Supplementary figure 1). Peak integration of size-exclusion chromatograms (SEC) showed high purity preparations (Supplementary figure 2). SDS-PAGE gels revealed a significant band between the reference ladders at 113 - 192 kDa (Figure 2a). The theoretical molecular mass of the spike is ∼136 kDa, suggesting that the distinct mobility reflects additional mass bound to the protein.

**Figure 2.**
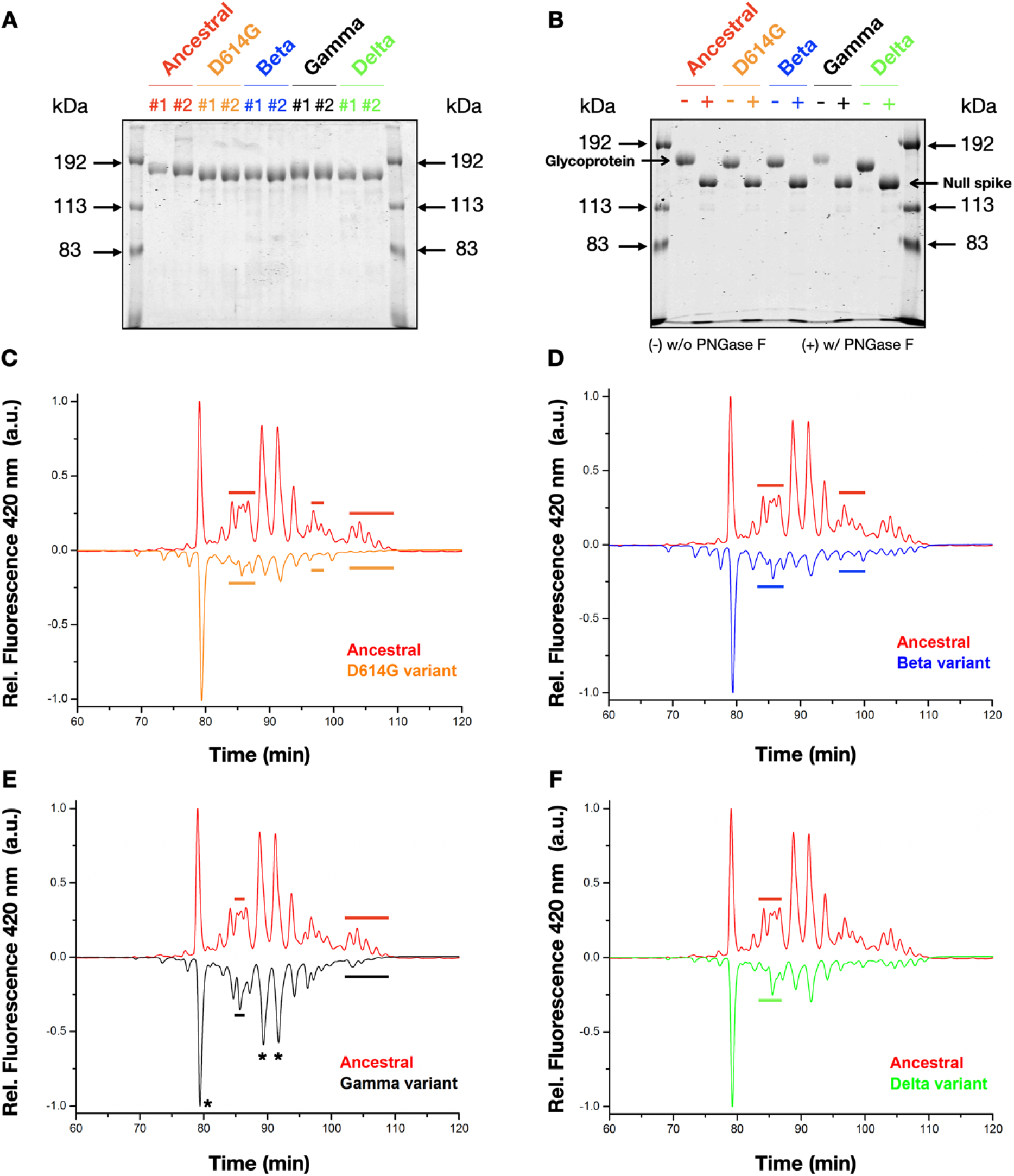
Spike protein glycosylation profile. **A, B**, Images of SDS-PAGE gels showing **(A)** two representative batches (#1 and #2) of purified spikes and **(B)** spikes in the absence (-) or in the presence (+) of PNGase F. Positions of the glycosylated and deglycosylated (“null”) spikes are shown in the gel. **C-F**, Line plots showing the relative fluorescence as a function of time from hydrophilic chromatography (HILIC) of **(C)** ancestral vs. D614G, **(D)** ancestral vs. beta, **(E)** ancestral vs. gamma, and **(F)** ancestral vs. delta. Results are flipped to facilitate visualization. Black asterisks show abundant glycan peaks. Thick horizontal bars show major differences relative to the ancestral spike.

**Figure 3.**
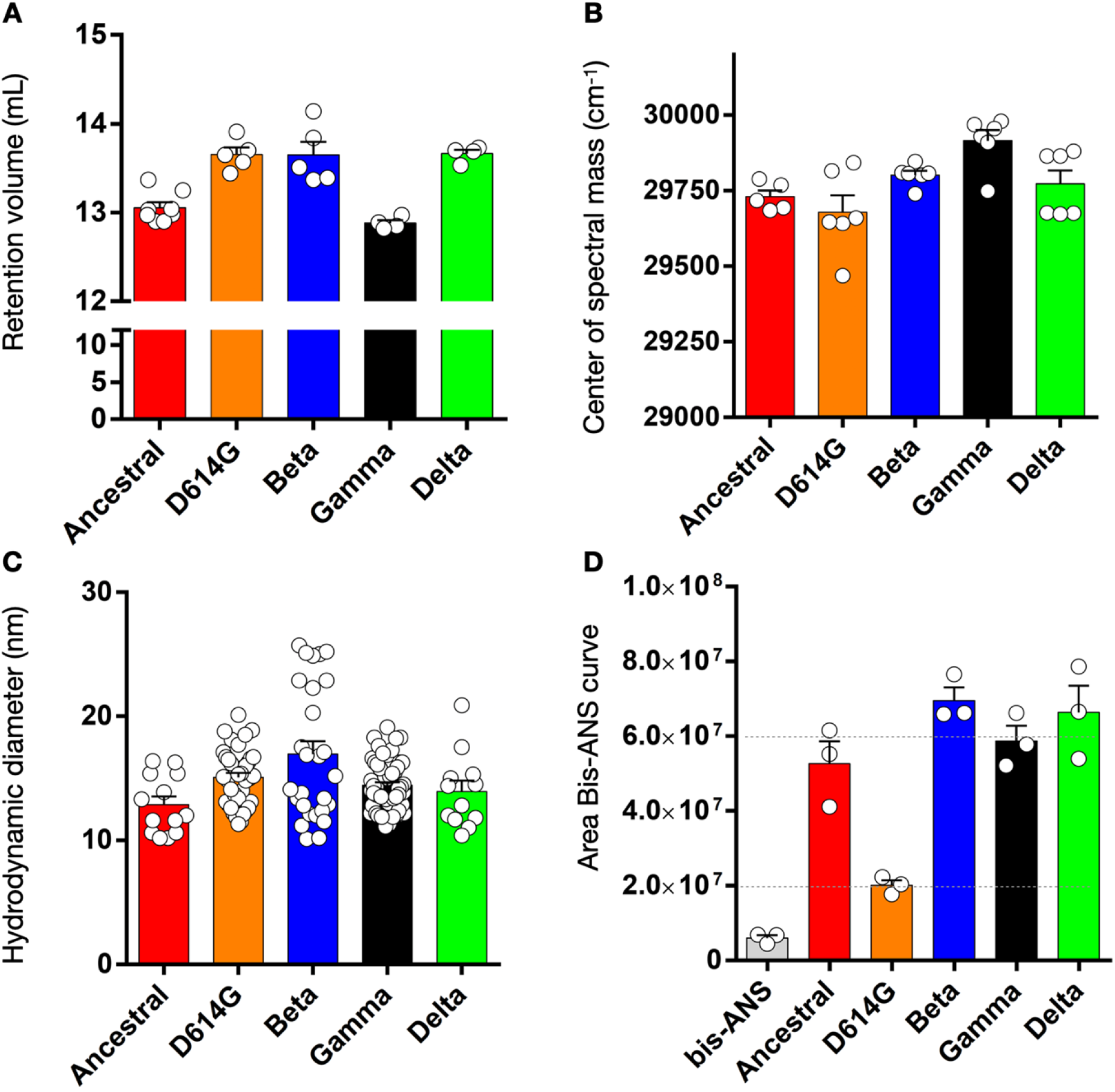
Structural features of SARS-CoV-2 spikes. **A-D**, Scatter plots showing **(A)** the retention volumes of studied spikes obtained from analytical size-exclusion chromatography. Data shows the avg. ± s.e.m. of independent protein batch preparations (n = number of data points for each series); **(B)** the center of spectral mass of studied spikes obtained from the emission fluorescence spectrum upon excitation at 280 nm. Data shows the avg. ± s.e.m. of at least three independent protein batches; **(C)** the hydrodynamic diameter of studied spikes determined by dynamic light scattering. Data is the avg. ± s.e.m. of several technical replicates from the same protein batch; **(D)** Area under the Bis-ANS curve of studied spikes obtained from the emission fluorescence spectrum upon excitation at 360 nm. Data shows the avg. ± s.e.m. of independent protein batches (n = 3). The gray bar shows bis-ANS fluorescence noise in the presence of buffer.

We noticed gentle mobility differences among purified variants. While the ancestral and gamma variants are slightly higher in the gels, the D614G, the beta, and the delta variants are lower (Figure 2a). Proteins on the surface of viruses are highly glycosylated, which also happens with the SARS-CoV-2 spike(31). We performed an enzyme-induced cleavage of potential N-linked glycans to show that the mobility shifts observed in the gels (Figure 2b) are clear-cut evidence that purified spikes are all in their glycosylated forms.

The gel positions of the ancestral and the gamma strains against the D614G, beta, and delta variants (Figure 2a) raise whether their glycan content is the same. Using hydrophilic chromatography (Figure 2c-f), we noticed that the change in the relative abundance of the three major glycan peaks is smaller between the ancestral and the gamma variant (Figure 2e, black asterisks) than when compared to the D614G, beta, and delta variants (Figure 2c, d, f). The result is indicative that the ancestral and the gamma have a higher content of glycans when compared to the others, and it goes in line with the slightly higher position of these proteins in the gel of figure 2a. In terms of peak profile, it is harder to identify pronounced differences, but some have been annotated by thick horizontal bars (Figure 2c-f).

### Structural features of SARS-CoV-2 spike variants

We performed biophysical experiments to comparatively understand variant differences and get insights into their structural behavior. To check whether purified proteins are forming trimers, we performed analytical SEC (Figure 3a and Supplementary figure 4a). All five proteins revealed a retention volume close to the reference marker at 670 kDa (Supplementary figure 4a). The theoretical mass of the trimer is 408 kDa, excluding any attached glycans. Thus, the results indicate a trimeric spike in solution.

Moreover, we noticed gentle differences in the retention values among studied variants (Supplementary figure 4a). To check the robustness of those differences, we produced additional batches of each variant. We consistently observed the differences in retention values among the variants (Figure 3a and Table 1). The ancestral strain and the gamma variant showed smaller retention values when compared to the D614G, beta, and delta (Figure 3a). Thus, the SEC data further supports the higher glycan content attached to the ancestral strain and the gamma variant (Figure 2a and e).

**Table 1.** Biophysical parameters of spike variants

|  | DLS |  | SEC |  | Fluorescence |  |  |
| --- | --- | --- | --- | --- | --- | --- | --- |
|  | D<sub>H</sub> (nm) | σ | Ret.<sub>vol</sub> (mL) | *T<sub>50%</sub> (min) | ν (cm<sup>-1</sup>) | **ΔG<sub>d</sub><sup>H<sub>2</sub>O</sup> (kcal/mol) | Bis-ANS (a.u.) |
| **Ancestral** | 12.8 ± 0.65 | 5.5 | 13 ± 0.05 | ~ 18 | 29,730 ± 20 | 1.33 | 52 ± 6 |
| **D614G** | 15 ± 0.36 | 5.0 | 13.6 ± 0.07 | n.d. | 29,678 ± 55 | 1.11 | 20 ± 1.3 |
| **Beta** | 16.9 ± 1 | 28.9 | 13.6 ± 0.14 | n.d. | 29,801 ± 14 | 0.88 | 69 ± 3.5 |
| **Gamma** | 14.4 ± 0.26 | 3.9 | 12.8 ± 0.03 | ~ 43 | 29,915 ± 34 | 1.09 | 58 ± 4 |
| **Delta** | 13.9 ± 0.87 | 9.1 | 13.6 ± 0.04 | n.d. | 29,772 ± 43 | 0.89 | 66 ± 7 |

\* Trimer-to-monomer spike dissociation at 40°C. (n.d. not detected)
\*\* Two-state approximation from folded trimers to unfolded monomers.

To explore whether surface differences and the altered polypeptide sequence would impact the structure of the variants, we evaluated the secondary and tertiary structure content by circular dichroism (Supplementary figure 4b) and intrinsic fluorescence spectroscopy (Figure 3b and Supplementary figures 5a, c, e, g, and i), respectively. Using dynamic light scattering (DLS) we measured the overall hydrodynamic spikes’ diameter (Figure 3c and Supplementary figures 6a-e). Finally, we used fluorescence to measure the surface exposure of the proteins’ hydrophobic motifs (Figure 3d and Supplementary figures 5b, d, f, h, and j). Far-UV circular dichroism spectra showed the expected behavior for an alpha-helix enriched protein with negative peaks at 222 and 208 nm and no differences among studied variants (Supplementary figure 4b). Fluorescence emission spectra upon excitation at 280 nm revealed λ_max_ of ca. 320 nm for all variants, but gentle differences in fluorescence intensity at λ_max_ (Supplementary figures 5a, c, e, g, i). The center of spectral mass (*ν*) is sensitive to inform whether excited probes (Tyr and Trp residues) are more exposed or hidden into the spikes’ structure. The smaller the value, the more exposed the Tyr/Trp residues are to the surface. No significant differences in *ν* values have been captured, indicating that the overall spikes’ structure is folded (Figure 3b and Table 1). Hydrodynamic diameter (D_H_) measurements revealed avg. ± s.e.m. values of 12.8 ± 0.65 nm (n = 13), 15 ± 0.36 nm (n = 39), 16.9 ± 1 nm (n = 27), 14.4 ± 0.26 nm (n = 55), and 13.9 ± 0.87 nm (n = 12) for the ancestral, D614G, beta, gamma, and delta, respectively (Figure 3c and Table 1). The beta strain showed the highest D_H_ variance (σ) value (σ = 28.9). The second highest value was for the delta strain (σ = 9.1), suggesting that these proteins show more heterogeneous structures in solution (Figure 3c and Table 1).

To learn about the content of hydrophobic motifs on the proteins’ surface, we measured the emission fluorescence spectra of a molecular probe (bis-ANS). Bis-ANS significantly increases its quantum yield upon binding to hydrophobic protein surfaces (Figure 3d and Supplementary figures 5b, d, f, h, and j). When bis-ANS is free in solution, the spectrum gives a λ_max_ of ca. 530-540 nm but undergoes a blue-shift to 470-490 nm and an intensity increment upon binding (Supplementary figure 5k). In contrast to all other spikes, the D614G presented a three-fold bis-ANS intensity reduction suggesting that the Asp-to-Gly substitution at position 614 can reduce the hydrophobic motifs’ surface exposure (Figure 3d and Table 1). It is worth mentioning that, except for the ancestral spike, all other studied variants have the D614G substitution, which made us conclude that the additional alterations of the beta, gamma, and delta could revert the lower hydrophobic surface exposure of the D614G variant.

### Modulating the structural stability of SARS-CoV-2 spike and variants

Because structural rigidity of surface antigens may have a role in transmissibility tendencies(32, 33), we decided to explore the stability of spike variants by using chemical and physical strategies. We first attempted to find conditions to discriminate energetic gaps related to conformational changes, dissociation, and unfolding events. We performed chemical-induced titrations with increasing concentrations of guanidine to challenge the spikes’ structure. We found buffer compositions that provided information about the abovementioned events using the ancestral spike. We noticed that a tris-based buffer vs. phosphate buffering leads to drastic changes in chemical-induced titrations (Figure 4a, b). While phosphate buffering showed two-state transition traces (Figure 4a), a tris-based buffer (see Methods) revealed an unexpected four-state transition (Figure 4b and Supplementary figures 7a-Aa). To check the consistency of the four-state transition, we produced additional protein batches, and all reproduced the four-state profile (Supplementary figure 7Bb). The reliability of these traces motivated us to explore the enriched species in each transition. In the first-transition range, the fluorescence emission spectra showed a gentle intensity increase followed by a smooth red-shift (Supplementary figures 7a-m, from 0 up to 0.3 M). The next transition was a delicate blue-shift accompanied by an intensity reduction (Supplementary figures 7n-t, from 0.4 to 1.9 M). Finally, we noticed a strong red-shift from λ_max_ at ca. 330 nm to 356 nm, possibly explained by an unfolding process in which the intrinsic probes moved away from hydrophobic moieties into the solvent (Supplementary figures 7u-Aa).

**Figure 4.**
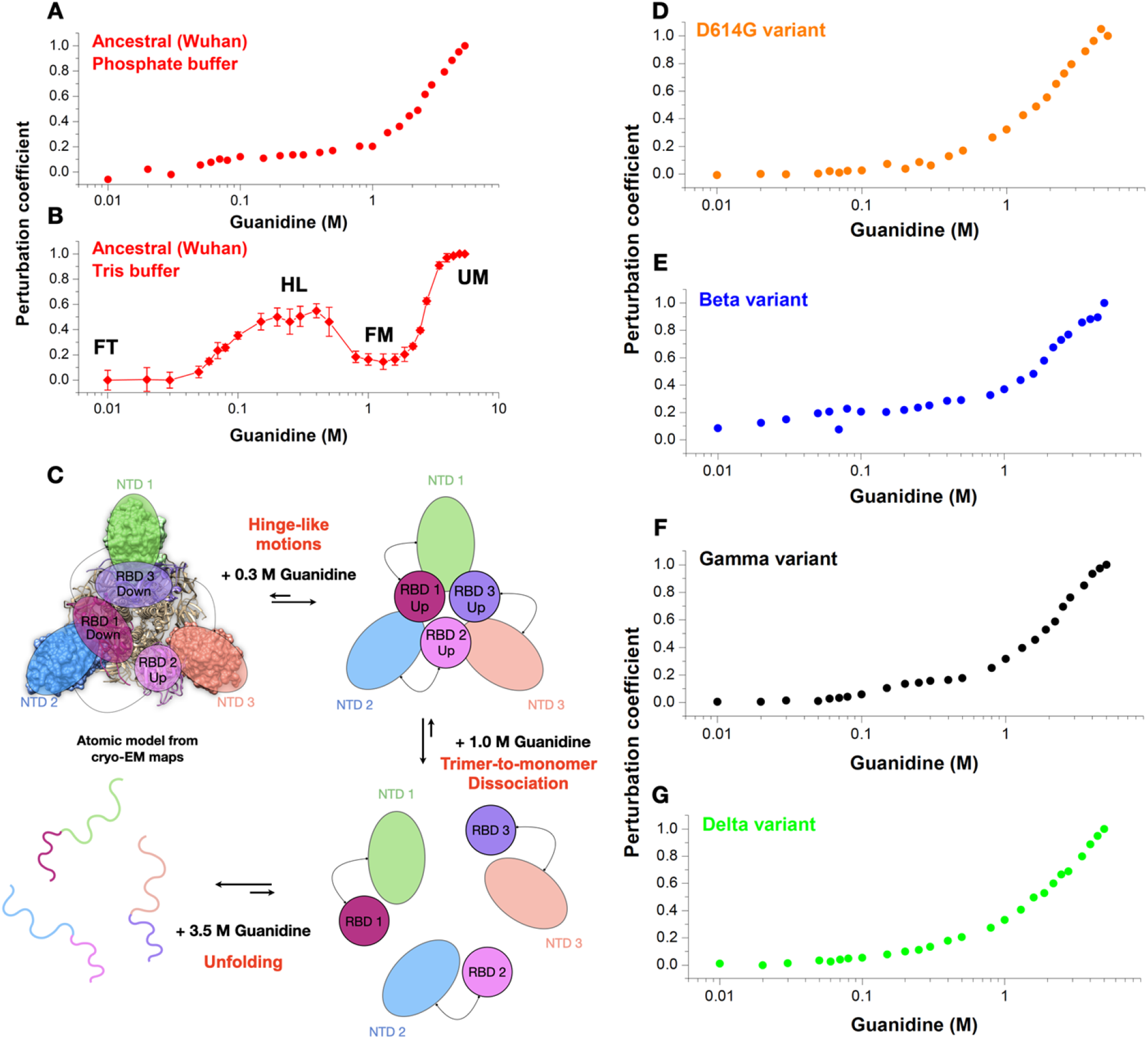
Structural stability of SARS-CoV-2 spikes. **A, B, D-G**, Dot plots showing the perturbation coefficient as a function of guanidine increments for the ancestral spike in the presence of **(A)** phosphate buffer (tagless protein) or **(B)** tris-based buffer (tagged protein, see Methods for details), **(D)** D614G, **(E)** beta, **(F)** gamma, and **(G)** delta variants. Curves corresponding to panels D-G were carried out in phosphate buffer and tagless proteins. Abbreviations on panel B stand for FT, folded trimer; HL, hinge-like; FM, folded monomer; UM, unfolded monomer. **C**, Schematics showing spike top views to illustrate the interpretation of the data. At 0.3 M of guanidine, the data supports RBD hinge-like motions. At this stage of knowledge, we cannot rule out whether all three RBDs are facing up. At 1 M of guanidine, the data indicates trimer-to-monomer dissociation. At concentrations higher than 3.5 M guanidine, monomer unfolding is achieved.

Using the ancestral strain, we performed analytical SEC and DLS at 0.3 and 1 M of guanidine to learn about the second and third plateau stages (Supplementary figures 8a and b). At 0.3 M of guanidine, the trimeric spike showed a larger retention volume when compared to the folded trimer (Supplementary figures 8a) and, in agreement with that, an amenable decrease in average D_H_ values (Supplementary figure 8b). The SEC and DLS data indicate a tight trimer at 0.3 M of guanidine that is reasonably interpreted as hinge-like motions occurring in the spikes’ RBD. An explanation to further supports our interpretation comes from the location of the intrinsic probes throughout the spikes’ structure (Supplementary figures 8c-e). Each protomer contains 53 Tyr (4.1%) and 12 Trp (0.9%) residues. Of those, 30 Tyr and four Trp residues are located within the RBD and the NTD (Supplementary figures 8c-e). Both domains comprise approximately a third of the spike sequence (1278 residues), meaning that the remaining 23 Tyr and eight Trp span across more than 750 residues.

Further, an RBD Tyr cluster is located within the RBD that rests in a position that allows partial occlusion when two RBDs are located down as in head-to-tail protection (Supplementary figure 8f). As RBD hinge-like motions occur and, at least, one RBD goes up, Tyr protection is unlocked, leading to their exposure to the solvent. This observation explains the red-shift within the 0 - 0.3 M guanidine range and the SEC and DLS, as hinge-like RBD motions reasonably favor a packed structure. At 1 M of guanidine, SEC and DLS revealed a minor peak consistent with a trimer-to-monomer dissociation and more considerable variance and D_H_ values pointing to the formation of more heterogeneous species in solution (Supplementary figures 8a and b). Additionally, chromatograms presented a full width at half maximum (FWHM) of 1.63, 1.55, and 2.98 for the spike in the absence and presence of 0.3 and 1 M of guanidine, respectively. These values indicate spike displaces from a narrow ensemble of species at 0.3 M to a broader ensemble at 1 M, consistent with an equilibrium shift to trimer dissociation. Figure 4c summarizes the proposed events occurring during the chemical-induced reaction recorded by fluorescence spectroscopy in Figure 4b.

Hereafter we focused on the folding-unfolding equilibrium to understand the structural rigidity of the variants. We performed chemical-induced reactions with phosphate in which the major detected species are folded trimers (FT) and unfolded monomers (UM). All curves revealed the two-state sigmoidal shape and amenable differences at the transition points (Figures 4d-g), reflecting the structural stability of the FT-UM reaction. Using a two-state model and linear regression of the transition points and thermodynamic equations (see Methods), we calculated the Gibbs free energy change in the absence of guanidine for the FT-UM reaction (ΔG _d_^H^_2_^O^) of each variant. The ΔG_d_^H^_2_^O^) values were 1.33, 1.11, 0.88, 1.09, and 0.89 kcal/mol for the ancestral, D614G, beta, gamma, and delta, respectively (Supplementary figures 9a-e and Table 1). This result indicates that spike tends to accumulate alterations that lead to less structural rigidity in virus adaptation. Using a tris-based buffer and the transition points related to the unfolding process, we figured out a ΔG _d_^H^_2_^O^) ^H O^ of 3.62 kcal/mol, which means a more stable structure under these conditions (Supplementary figure 9f).

### Modulating SARS-CoV-2 spike by physical forces

We next designed experiments to understand the trimer-to-monomer equilibrium of each variant. We captured their dissociation tendencies by increasing the incubation time to 40°C, followed by SEC injections at room temperature (Figure 5a-e). For the ancestral strain and the gamma variant, the peak corresponding to the trimeric spike (T) decreased concomitantly to the increase of the spikes’ monomeric peak (M) (Figures 5a and b). The D614G variant has not shown any traces of trimer dissociation within the evaluated timeframe (Figure 5c). In contrast, the beta and delta variants revealed smaller amounts of monomers resulting in a shoulder-like peak (Figures 5d and e). The peak areas of the trimers and monomers as a function of time showed crossover points for the ancestral and the gamma variants (Figure 5f, red and black plots). The crossover point indicates the T_50%_ when the reaction has approximately 50% of trimers and monomers in the solution. The T_50%_ values were ca. 18 min and 43 min for the ancestral and the gamma, respectively.

**Figure 5.**
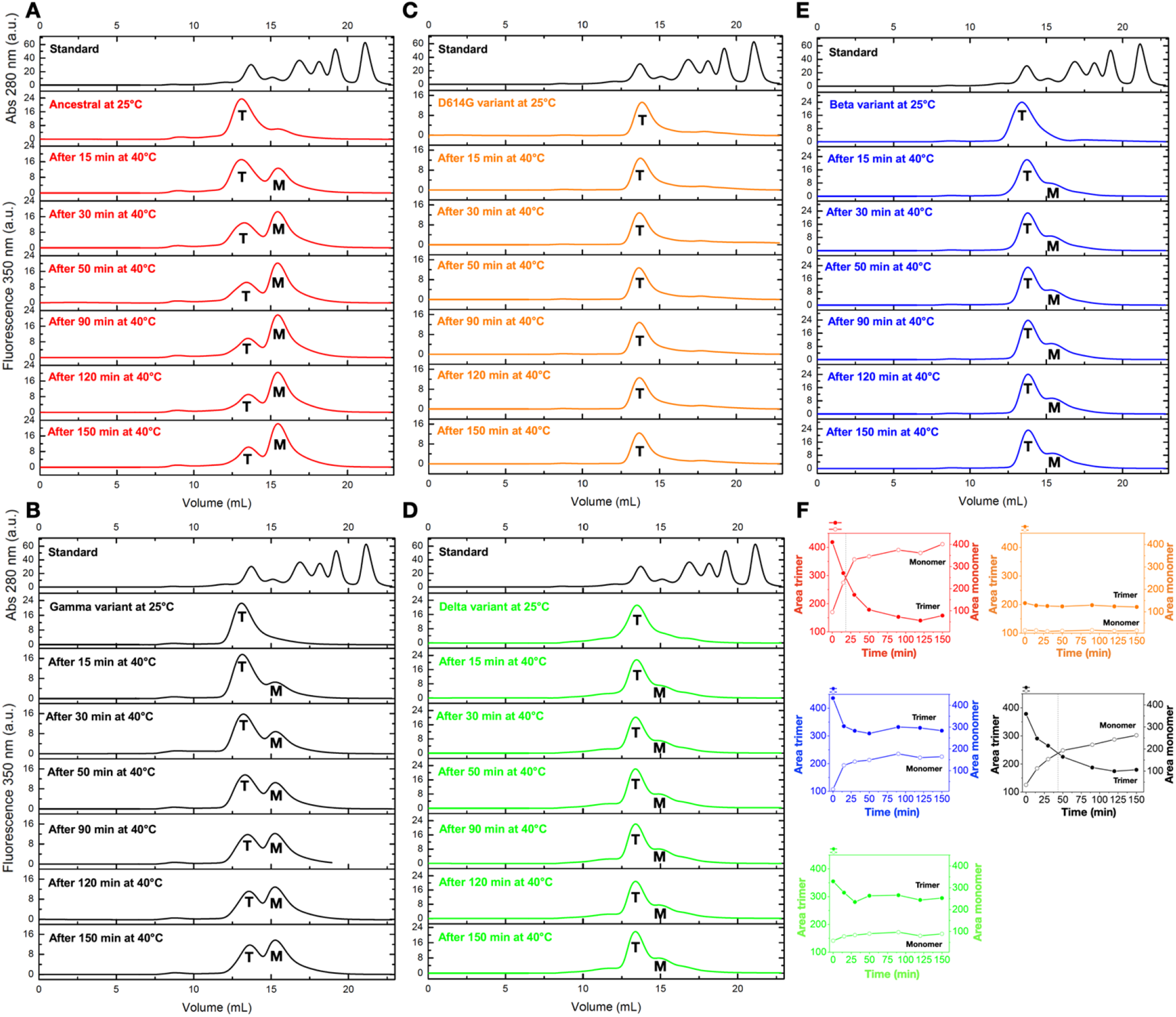
Trimer-to-monomer spike dissociation. **A-E**, Collection of line plots showing the fluorescence signal at 350 nm as a function of retention volume for **(A)** the ancestral strain in red, **(B)** gamma in black, **(C)** D614G in orange, **(D)** delta in green, and **(E)** beta in blue at 25°C and after several times at 40°C. T and M stand for trimer and monomer, respectively. The absorbance at 280 nm of the calibration standard is shown at the top of each experiment. **F**, Double-Y plots show the integration area corresponding to the trimers (filled dots) and the monomers (empty dots). The color code is the same as for panels A-E. Crossover points (T_50%_) are highlighted by vertical dashed lines for the ancestral strain (red) and gamma variant (black).

We also evaluated the effects of hydrostatic pressure as a physical force acting primarily on the volumetric properties of macromolecules(34). By following the bis-ANS fluorescence as a function of pressure, we noticed that the delta variant is more susceptible to pressure-induced hydrophobic exposure when compared to the others (Figure 6a and Supplementary figure 10a-e). In terms of light scattering intensity, all spike versions showed a similar tendency to decrease approximately 30% of the light scattering intensity against pressure increments, except for the beta variant, which showed a lower decrease as a function of the pressure exerted (Figure 6b and Supplementary figure 10f-j). After returning to atmospheric pressure, light scattering values returned partially (not shown). Further, no retention volume changes have been noticed when depressurized samples have been injected into analytical SEC (not shown). The results are consistent with a reversible trimer dissociation, with the delta showing the highest susceptibility of hydrophobic exposure during dissociation (Figure 6a) and the beta showing the lesser (Figure 6b).

**Figure 6.**
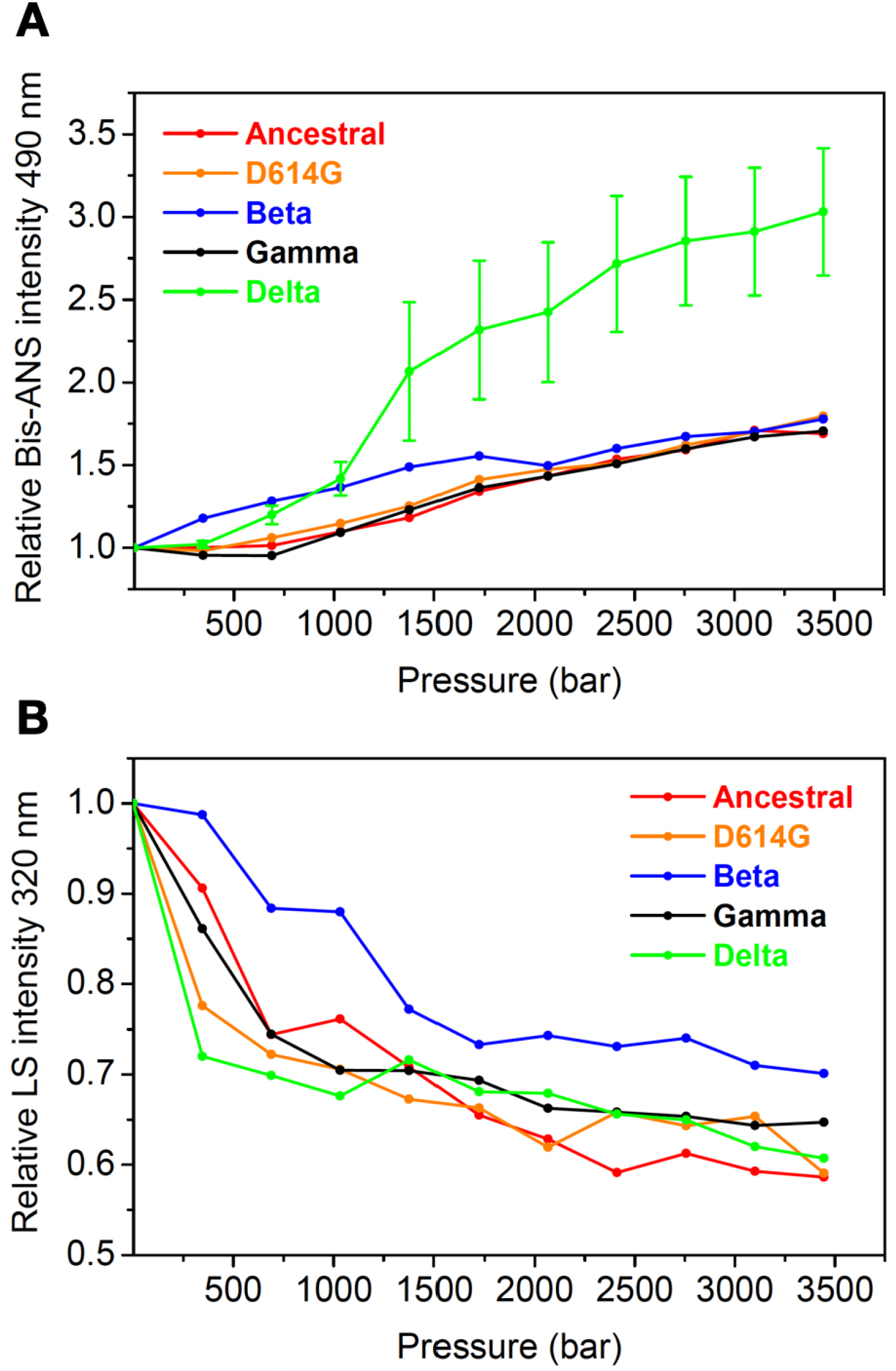
Pressure-induced trimer-to-monomer spike dissociation. **(A)** Relative bis-ANS fluorescence intensity and **(B)** light scattering intensity as a function of pressure increments for studied spikes. Data is shown as the avg. ± s.e.m. of independent experiments with three delta variant batches (n = 3). All other variants show n =1 as data is highly reproducible.

## Discussion

The data revealed features between the gamma spike and the ancestral spike. They present equal mobility shifts on SDS-PAGE gels because of their similar glycan abundance (Figure 2). Moreover, they share similar retention volumes (Figure 3a) and a detectable pattern of trimer-to-monomer dissociation despite different T_50%_ values (Figures 5a and b). These features are unique to the ancestral and gamma spikes and not observed for the D614G, beta, and delta variants. The ancestral, the D614G, and the gamma revealed a higher affinity to ACE-2 than the beta and the delta (Supplementary figure 3). The D614G variant showed unique features such as a 3-fold reduction in bis-ANS fluorescence (Figure 3d) and no trimer dissociation after 150 minutes at 40°C (Figure 5c). Further, the beta variant showed the highest variance in D_H_ values, followed by the delta (Figure 3c).

Using chemical-induced unfolding, we provide evidence to argue that during SARS-CoV-2 adaptation, the spike glycoprotein accumulates changes that lead to less structural rigidity (Figure 4). *In silico* data has shown that SARS-CoV-2 was selected for amino acid substitutions, resulting in more stable spikes(35). Still, a significant drawback of this study is the absence of glycosylation effects on protein stability(35). Our results point opposite, and data is provided using glycosylated spikes. We also captured energetic gaps for the ancestral spike that is reasonably due to RBD hinge-like motions, trimer dissociation, and unfolding (Figure 4b and c). The sequential events of conformational change and trimer dissociation in spike unfolding open up insights into understanding more general protein folding mechanisms in complex biological systems. However, we cannot distinguish whether RBDs adopt the canonical up positioning required for receptor binding.

The features of D614G are intriguing. D614G-containing viruses replaced the ancestral strain to reach worldwide near-fixation within a few months after the COVID-19 pandemic started(14). This substitution became dominant by April 2020(36) and appeared to be a fitness advantage rather than genetic drift as its frequency increased globally and during co-circulation within individual regions(15). SARS-CoV-2 carrying G614 enhances viral replication in human lung cells, expanding the infectivity and stability of virions(15). Further, G614-infected hamsters produced higher infectious titers in nasal washes and the trachea, supporting higher viral loads in the upper respiratory tract of COVID-19 patients(15, 37). Mechanistically, the D614 interactions are a matter of debate still. Structures from April 2020 revealed that the D614 of one protomer forms an H-bond with the T859 at the S2 of the adjacent protomer(7, 38). However, structures from August 2020 revealed that D614 contacts K854, forming a salt bridge(39) (Figure 1b). G614 provides a shorter side chain abolishing interprotomer interaction. The consequence is a more dilated packing and a range of open conformations, presumably because of increased mainchain flexibility and local unfolding. Cryo-EM studies on D614 (ancestral) and G614 spikes revealed a dramatic flip in the ratio of open to closed conformations. Particle distributions indicated a shift from 82% closed and 18% open for D614 to 42% closed and 58% open for G614, with a substantial fraction presenting two-open RBDs (39%) and all-open RBDs (18%)(40). The D614G structural flexibility seems to be related to its higher infectivity, and G614 persists in all subsequent variants that emerged. However, we observed features of D614G that are not shared among the studied VOCs, which made us conclude that the other mutations the virus accumulated had different implications. Hence, the D614G mutation alone was a relevant aspect of the virus spreading at the early stages of the pandemic. Compared with the ancestral, beta, gamma, and delta strains, the D614G variant was the only one to present a substantial reduction in surface exposure of hydrophobic motifs (3-fold) and trimer persistence after two hours under mid-temperature incubation (40°C).

Several hypotheses attempted to explain the effects of D614G for infectivity and transmissibility. One is that the protomer impairment would cause local unfolding and prime the fusion peptide to a fusion-prone conformation(41, 42). Another possibility is that local unfolding may trigger the exposure of asparagine sites amenable to the glycosylation machinery(43). Glycan modification has been shown to modulate spike conformation and host cell invasion(44, 45). Recently, N-glycan at position N343 revealed a gating role facilitating RBD opening(45), and surprisingly, D614G has shown substantial differences in glycan content at N343 compared with the WA1 ancestral strain(43). This finding aligns with our HILIC profile that shows differences between the ancestral and D614G variant (Figure 2c). The combinatory events of less hydrophobic surface exposure (Figure 3d) and trimer persistence (Figure 5c) with D614G populating more open and flexible conformations(40, 41) support a model of compensatory attributes (screening flexibility vs. higher immunogenicity) for fitness advantage. It is worth mentioning that D614G reduced ACE2 affinity due to a faster dissociation rate(40), supporting one scenario of the screening flexibility model. In our hands, the ancestral, the D614G, and the gamma revealed higher ACE-2 affinity when compared to the beta and delta spikes as measured by EC_50_ values (Supplementary figure 3). The lesser the content of D614G hydrophobic exposure, the weaker the antigen stifness is to generate a more structurally adaptable antigen with unlocked RBD conformations. This phenomenon would help on two sides: screening more rapidly for potential ACE-2 interacting surfaces and binding more tightly once required. This interpretation reconciles apparent controversial results on D614G affinity for ACE-2.

Suppose equilibrium is displaced to all RBDs achieving a closed state. In that case, the RBD-ACE2 recognition motif is blocked, implying a strategy for the virus to protect the critical-binding site from the host immune response. Contrastingly, when RBD heterogeneity arises, viruses become susceptible to neutralizing antibodies that commonly target the RBD(14, 15, 37). D614G is probably a trade-off between a structurally adaptable antigen promoting a more significant advantage for ACE-2 screening at the cost of higher immunogenicity. Although D614G substitution persisted in subsequent variants, the beta, gamma, and delta reversed the lesser hydrophobic exposure (Figure 3d), presumably due to the acquisition of other alterations (Figure 1d-f). Altogether, we conclude that the decreased hydrophobic exposure and higher flexibility of D614G were critical for the virion to achieve fast worldwide near-fixation but irrelevant for long-term virus adaptation.

Hydrophobic interactions make a dominant contribution to protein stability(46). Less hydrophobic motifs at the surface mean a more significant contribution of hydrophobic interactions inward of the protein manifold, possibly tuning the structural stifness and flexibility ratio. Higher flexibility and lower surface hydrophobicity were observed in other unrelated molecules, including crystallin, heat repeat proteins(47, 48), and carbon nanotubes(49). This relation is explained by observing how the chemical topology of proteins impacts the water-density fluctuations of protein’s hydration shells, which is an adequate measure of hydrophobicity(50). Less hydrophobic surface exposure means that water is weakly repelled, which may help create a relaxed topology for heterogeneous ensembles and multiple states of openness.

Moreover, the D614G is slightly less stable than ancestral with ΔΔG_d_ of 0.22 kcal/mol, but equally stable when compared to the gamma (ΔΔG_d_ = 0.02 kcal/mol) (Table 1). All three variants (D614G, ancestral, and gamma) revealed similar spike stabilities and higher ACE-2 affinity than the beta and the delta, which points to some degree of RBD rearrangement and topological flexibility for ACE-2 binding. This interpretation is supported by weakened trimeric stability for the ancestral and the gamma (Figure 5) and D614G showing a 3-fold reduction in hydrophobic surface exposure (Figure 3d) and higher thermostability (Figure 5). The flexible nature of D614G differs from that observed for the ancestral and gamma spikes. D614G flexibility is accompanied by higher thermostability, likely resulting from an entropic compensation(51) due to rearrangement of hydrophobic interactions. In contrast, the ancestral and gamma flexibility results from a less stable trimeric arrangement.

The relation between structural rigidity of surface glycoproteins and their antigenicity goes beyond SARS-CoV-2. For example, influenza A undergoes constant antigenic evolution(32) while measles virus (MeV) exhibits homotypic antigenicity(33). In contrast to influenza A, the lower tolerance of MeV to gene diversity is associated with more rigid hemagglutinins. This observation explains the restricted repertoire of seasonal MeV strains and the effectiveness of measles vaccines for more extended periods(32, 33). Therefore, including a more flexible recombinantly-expressed D614G and gamma spike in COVID-19 vaccines seems a wise strategy to efficiently elicit host immune response against SARS-CoV-2.

The ancestral and the gamma spikes share similar retention volumes (Figure 3a) and equal mobility shifts on SDS-PAGE (Figure 2a) primarily due to corresponding glycan abundance (Figure 2e). They also have a discernible trimer-to-monomer dissociation pattern despite different T_50%_ values (Figures 5a and b). These shared properties are significant because they are not observed for D614G, beta, and delta variants. Recent biochemical results revealed that the gamma protein eluted in three distinct peaks, corresponding to the prefusion trimer, postfusion trimer, and dissociated monomers, with the prefusion trimer corresponding to less than 40% of the total protein, similarly to the ancestral protein(28, 52). The authors concluded that the trimer is not stable for this variant and raised uncertainty of why the gamma trimer dissociates(28). In our hands, the gamma and the ancestral trimer are eluted as a single peak, probably related to the mutations used in our gene construct to stabilize the protein trimer in the prefusion conformation^35^. However, we detected a trimer-to-monomer dissociation profile for both spikes (and not for the other studied variants), which is in line with the reported trimer instability for the ancestral strain and the gamma variant^27,50^.

Notwithstanding, the gamma trimer is more stable than the ancestral trimers, with T_50%_ values of 43 and 18 minutes, respectively. We believe the corresponding abundance of glycans bound to the surface of the ancestral and the gamma may exert, to some extent, a role in their trimer dissociation profile. It is worth mentioning that despite providing structural shielding, glycans are highly hydrophilic molecules, attracting water to their boundaries, which may contribute entropically to nucleate local dissociation at mid-temperatures. Alternatively, the presence of H655Y at the S2 would contribute to destabilizing the gamma trimer(28).

Observing distinct traces during chemical-induced unfolding (tris-based buffer vs. phosphate) does not mean that hinge-like and dissociation events vanish when reactions are performed with phosphate (Figure 4). While in the phosphate buffer, protein-solvent interactions presumably favor microscopic barriers not possible to be annotated by steady-state fluorescence, the tris-based composition is probably acting on stabilizing forces to allow the discrimination of energetic barriers. The distinction of each step on the way to unfolding is relevant to understanding more general mechanisms of protein unfolding and association in complex systems. The data supports a sequential unfolding model in which trimer dissociation is preceded by conformational changes to unlock quaternary RBD interactions. Following this, protomer unfolding happens cooperatively. It is worth mentioning that experiments are under equilibrium and represent steady-state measures at each guanidine concentration. Thus, plateau states are likely explained by a population-weighted average of multiple conformers enriched and trapped in solution and not by a single conformer.

In summary, there is an utmost necessity to explore the dynamics of the spike variants and the way conformational changes, glycosylation, and trimer stability would affect the ensemble of spike conformations and the status of up/down RBD positioning. For example, the Folding@home platform has contributed tremendously with 0.1 seconds of simulation data(53). This study predicts spike has a trade-off between making ACE-2 binding interfaces accessible and masking epitopes to circumvent immune response(53). Another study used single-molecule fluorescence resonance energy transfer (smFRET) to observe spike dynamics on virus particles. The authors found that spike samples four distinct conformational states forming an intermediate during ACE-2 recognition(54). Finally, hydrogen-deuterium exchange coupled with mass spectrometry revealed a spike conformation that interconverts slowly with the prefusion state and is more likely explained by an open trimer that exposes the S2 trimer interface(12). We experimentally compared the ancestral, D614G, beta, gamma, and delta variants to uncover their trimer stability and structural rigidity and understand spike evolution during the pandemic. Studies on the energetic landscape of the full-length spike variants in its glycosylated form are sparse but *sine qua non* to understand the antigenicity of circulating strains and help develop more effective vaccines.

## Methods

### SARS-CoV-2 spike protein constructs

The Wuhan ancestral gene construct (GenBank MN908947) encodes the ectodomain (aminoacids 1-1208) spike protein in the trimeric form and the prefusion conformation^35^. A pαH vector comprising this gene was kindly provided by B. Graham, VRC/NIH, and is also available from BEI Resources under #NR-52563. The sequence contains proline substitutions at residues 986 and 987, a GSAS substitution at the furin cleavage site (residues 682-685), a C-terminal T4 fibritin trimerization motif, an HRV-3C protease cleavage site, a TwinStrepTag®, and an 8xHis tag. Since the pαH vector does not have a cassette for selecting stable transfectants, it must be co-transfected with an empty vector containing the neomycin phosphotransferase selectable marker^51^. The genes coding for the D614G, beta, gamma, and delta variants were synthesized at Genscript (Piscataway, USA) and cloned into the pCIneo vector (Promega, USA) to facilitate stable expression in mammalian cells. The variants contained the respective mutations as follows

1. D614G: Asp-to-Gly substitution at position 614 (D614G)
2. Beta (lineage 20H or B.1.351): D80A, D215G, 242-244del, K417N, E484K, N501Y, D614G, A701V
3. Gamma (lineage 20J or P.1): L18F, T20N, P26S, D138Y, R190S, K417T, E484K, N501Y, D614G, H655Y, T1027I, V1176F
4. Delta (lineage 21A or B.1.617.2): T19R, G142D, 156-157del, R158G, L452R, T478K, D614G, P681R, D950N

### Production and purification of spike variants

Stable, recombinant HEK293 cell lines were generated to produce each spike protein by transfecting the suspension-adapted HEK293-3F6 (NRC, Canada) parental cell line with each plasmid described above. After the selection of stable transfectants, the resulting recombinant cell pools were cultured in HEK-TF medium (Xell, Germany) in a CO_2_ incubator at 37°C and 5% CO_2_ using shake flasks under orbital agitation (140 rpm, 2.5-cm stroke) or in 1.5-L stirred-tank bioreactors at setpoints of pH, temperature and dissolved oxygen of 7.1, 37°C and 40% of air saturation, respectively (EzControl, Applikon, The Netherlands). Cell concentration and viability were determined by trypan blue exclusion using an automated cell counter (Vi-Cell, Beckman Coulter), whereas glucose and lactate concentrations were monitored using a metabolite analyzer (YSI 2700, Yellow Springs Instruments). Spot blots determined the presence of spike protein in the supernatants as described earlier^51^. Cell-free supernatant was obtained by microfiltration with 0.45-µm PVDF membranes and injected into a 5-mL StrepTrap XT affinity chromatography column (Cytiva, Sweden) following the manufacturers’ instructions using an Äkta Purifier system (Cytiva, Sweden). The purification tag was removed using HRV-3C protease (Thermofisher, #88946) to yield the tagless protein unless otherwise stated. Protein concentration, purity, and identity in the eluted fractions were confirmed by NanoDrop (Thermofisher USA), SDS-PAGE, and Western blot analyses, respectively. An extinction coefficient of 135,845 M^-1^.cm^-1^ and a molecular mass of 136,908.9 Da (monomer of the polypeptide chain) was used for the Nanodrop setting. The purified protein obtained in the affinity chromatography was stored at -80°C and thawed when needed for experiments.

### SDS-PAGE of protein samples

We evaluated the reactions by 7% SDS-PAGE gels stained with Coomassie brilliant blue R250. We digitized the gels using the Odyssey scanner and software (LI-COR Biosciences). In selected samples indicated in Figure 2b, we carried out PNGase F digestion following the manufacturers’ instructions (New England BioLabs, NEB # P0704S). Control samples followed the same procedure except for the enzyme.

### Hydrophilic interaction liquid chromatography (HILIC-HPLC)

First, we digested spike samples with PNGase F using a commercial kit (New England BioLabs, NEB # P0704S). We adopted a modified protocol as follows: a sample volume of 40 μL at 0.5 mg/mL (20 μg of spike protein) was mixed with 10 μL of denaturing buffer (part of the kit, “10X”) and incubated for 10 min at 100°C. After cooling down, 10 μL of NP-40 solution, 10 μL of GlycoBuffer, and 1 μL of the PNGase F enzyme (all kit components) were added and incubated at 37°C for 2 hours. After that, we separated the free glycans by precipitating the remaining proteins with 300 μL of cold ethanol for 15 min at -20°C. The tubes were centrifuged, and the supernatants of each sample were collected. The precipitation step was repeated twice, and a pool of three supernatants collected per sample was dried in a vacuum concentrator (Eppendorf, Germany). To label glycans with a fluorophore, each dried sample was mixed with 5 μL of a solution containing 0.048 g/mL of 2-aminobenzamide (2-AB) and incubated at 65°C for 2 hours. 2-AB-labeled glycans and unbound 2-AB were separated by filter paper chromatography (Whatman) using acetonitrile. 2-AB-glycans were recovered from cropped filter papers after washing them with 500 μL of ultrapure water for 10 minutes. We performed the washing procedure four times to produce a final volume of 2 mL (4 × 500 μL). Samples were filtered with 0.45-μm PVDF membranes and dried overnight using a vacuum concentrator (Eppendorf, Germany).

Dried samples were resuspended with 40 μL of ultrapure water and 160 μL of acetonitrile. Half of this volume was directly injected at 0.4 mL/min into an HPLC system (Shimadzu, Japan) fitted with a TSK gel amide 4.6 mm x 25 cm, 5 μm column (Tosoh Bioscience) previously equilibrated with 20% of 50 mM ammonium formate pH 4.4 and 80% of acetonitrile HPLC grade (initial step). Glycans were recorded by fluorescence at 420 nm upon excitation at 330 nm using an RF-10A XL fluorescence detector from Shimadzu. Elution was accomplished using a linear gradient in which the initial step achieved 53% of 50 mM ammonium formate pH 4.4 and 47% of acetonitrile after 132 minutes.

### Mass spectrometry sample preparation

Each spike variant (25 μg) was mixed with an equivalent volume of 10% sodium dodecyl sulfate (SDS, USB #75819) and 100 mM triethylammonium bicarbonate buffer, pH 8.5 (TEAB, Sigma #T7408). Disulfide bonds were reduced by incubating the samples at 90°C for 10 minutes with dithiothreitol at a final concentration of 10 mM (DTT, Bio-Rad #161-0611), followed by incubation at 37°C for 1 hour. Samples were then incubated with iodoacetamide at a final concentration of 40 mM (IAA, Bio-Rad #163-2109) at room temperature in the dark for 30 minutes to alkylate disulfide bonds.

Phosphoric acid at 12% (Millipore, EMD #1.00264.1000) was added until each sample’s final concentration of 1.2%. Then, 350 μL of binding/washing buffer (100 mM TEAB in 90% methanol, Tedia #MS1922-001) was added, and the total volume was loaded onto suspension trapping (S-Trap) columns (Protifi, #C02-mini-80). S-Trap protocol was performed following the S-Trap manufacturer’s instructions. Trapped proteins were incubated with PNGase-F (New England BioLabs, NEB # P0704S) (20 units/μg of protein) at 37 °C for 18h. PNGase F was previously diluted with 50 mM TEAB until a final volume of 25 μL. Tryptic (Promega, #V511A) (E:S, 1:50 w/w) digestion was conducted at 37°C for 16 hours. The enzyme was diluted in 50 mM TEAB until a final volume of 100 μL. Tryptic peptides were dried in a vacuum concentrator (Martin Christ, Germany) and resuspended in formic acid 0.1%. Peptides were quantified using QuBit Assay (Thermo Scientific, #Q33212), following the manufacturer’s instructions.

### Reverse phase nano-liquid chromatography tandem mass spectrometry and data analysis

Tryptic peptides (2 μg) were analyzed in an EASY-1000 nLC system (Thermo Scientific # LC120) coupled to a Q-Exactive Plus mass spectrometer (Thermo Scientific). The sample was loaded in a trap-column Acclaim PepMap™ 75µm x 2cm, nanoViper C18, 3µm, 100Å (Thermo Scientific, #164946) and separated in an analytical EASY-Spray™ Column PepMap™ Column (75µm x 25cm, C18, 2µm, 100Å, Thermo Scientific, # 03-251-871). The mobile phases were 5% acetonitrile/0.1% formic acid (Solvent A) and 95% acetonitrile /0.1% formic acid (Solvent B). The gradient used was 10-45% B for 40 min, 45-70% B for 8 min, 70-95% for 5 min, and 95% for 7 min, at 300 nL/min of flow rate.

The mass spectrometry analysis employed the full-MS/DDA-MS2 and positive polarity modes. A dynamic exclusion list of 45 s and spray voltage at 1.9 kV was set for all studies. The parameter settings for the full scan were 1 microscan, 70,000 resolution at m/z 200, AGC target of 3E6 ions, 100 ms maximum injection time, and the range of mass acquired was m/z 350 to 2000 m/z. The top 20 DDA-MS2 parameters were 17,500 resolution, m/z 200, AGC target of 1E6 ions, 50 ms maximum injection time, m/z 2.0 of isolation window, minimum intensity threshold of 2E5 ions equipped with High-energy Collision Dissociation (HCD) cell using a normalized collision energy of 30 NCE.

The raw data were imported to Proteome Discoverer 2.4 (Thermofisher Scientific) software to perform protein identification analysis. A homemade database was created, including three sequences of spike variants (D614G, Gamma/P.1, and Delta/B.1.617.2). Additionally, a human database from UniProt (reviewed database with canonical and isoforms, June 2021) was used. Two searches were executed with total and semi-tryptic peptides, two missed cleavages were considered. Carbamidomethylation (C) was set as a fixed modification, whereas methionine oxidation, asparagine deamidation, and acetylation (protein N-terminal) were selected as variable modifications. The mass tolerance for precursor ions was 10 ppm and 0.1 Da for the fragment ions. The False Discovery Rate (FDR) was fixed for protein and peptide validation with a cutoff score minor to 1% at the protein, peptide and PSM levels. Proteins were grouped in master proteins using the maximum parsimony principle.

### Analytic size-exclusion chromatography (A-SEC)

Spike variants at 25 μg/mL (injection volume: 250 μL) were directly injected into a Superose 6 increase 10/300 GL (Cytiva, #29091596). All runs were performed in phosphate buffer saline, PBS (140 mM NaCl, 2.8 mM KCl, 10 mM Na_2_HPO_4_, and 1.8 mM KH_2_PO_4_ pH 7.4) at 0.7 mL/min. Fluorescence was recorded at 350 nm upon excitation at 280 nm using a high-performance liquid chromatography system (Shimadzu Inc.). The column was calibrated using A_280nm_ with thyroglobulin (670 kDa), γ-globulin (158 kDa), ovalbumin (44 kDa), myoglobin (17 kDa), and vitamin B12 (1.35 kDa) (Bio-Rad, #151-1901). Column void was determined using blue dextran (2,000 kDa) at A_380nm_ and A_620nm_ (Merck, #D5751).

For trimer-to-monomer dissociation studies, a water bath (Fisher Scientific) at 40°C was kept close to the HPLC system, and samples were injected immediately after each time point (Figure 5). All HPLC runs were performed at room temperature. For guanidine experiments (Supplementary figure 7), all runs were performed with the respective guanidine concentration (0.3 or 1 M).

### Human ACE2 affinity

An indirect ELISA was performed to measure the affinity of the full-length ancestral, D614G, beta, gamma, and delta spike glycoprotein against the ACE-2 Fc protein. First, protein quantification was achieved by Pierce™ BCA Protein Assay Kit (Thermo Scientific, Cat# 23227, Batch# VH311372). 96-well plates were coated overnight at 4°C with 3-fold dilutions of each spike glycoprotein variant (tested concentrations between 50 μg/ml and 23 ng/ml). The 96-well plates were then washed with 0.05% PBS-Tween 20 (PBST) and blocked with 1% BSA in PBS (Sigma-Aldrich, Cat# A9647-50G, Lot# SLCH5268). Next, the plates were incubated with 2 μg/ml ACE-2 Fc protein (GenScript, Cat# Z03484-1, Lot# B2011010) at 37°C for 30 min and washed three times with 0.05% PBST. After that, anti-human IgG (Fc specific)-peroxidase antibody (Sigma-Aldrich, Cat# SAB3701282-2MG, Lot# RI34900) was added at 1:10,000 dilution, and the plate was incubated at room temperature for 1 h. After rewashing the plate, TMB substrate (Life Technologies, Cat# 2023, Lot# 11323202-7) was added, and the reaction was stopped after 20 min at room temperature with HCl 1 N. The absorbance was read at 450 nm with 655 nm background compensation in iMark Microplate Absorbance Reader (BIO-RAD Laboratories). The half-maximal effective response (EC_50_) was carried out by fitting the curve to a four-parameter logistic regression by GraphPad Prism version 8 (GraphPad Software, San Diego, California, USA).

### Circular dichroism (CD)

Far-UV spectra were recorded in a ChiraScan spectropolarimeter (Applied Photophysics) at wavelengths ranging from 200 to 260 nm at 25°C and 0.4 nm step size. Data is the average of five accumulation scans (Supplementary figure 3b). Raw ellipticities [θ] were recorded at a protein concentration of 50 μg/mL (final volume 300 μL) in a 2 mm path length cuvette using PBS. Data is shown as the mean residue ellipticity [Φ;] and calculated as follows

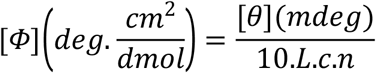

where *L* is the path length in centimeters, *c* is the molar concentration, and *n* is the number of peptide bonds (number of amino acids - 1).

### Steady-state fluorescence spectroscopy

All experiments were performed using spike variants at 20 μg/mL. Fluorescence emission measurements were acquired using an ISSK2 spectrofluorometer (ISS Inc.) equipped with a high-pressure cell (ISS Inc.). Samples were excited at 280 nm to measure Tyr + Trp, and emission was recorded from 300 to 400 nm. Changes in the fluorescence spectra for each spike variant was quantified as the center of spectral mass (*ν*) (equation 1), where *Fi* stands for the emitted fluorescence at wave number λ.

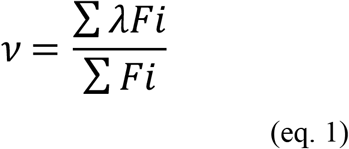

We used 4,4’-dianilino-1,1’-binaphthyl-5,5’-disulfonic acid, bis-ANS to measure hydrophobic surface exposure (Invitrogen, #B153). A bis-ANS stock solution in phosphate buffer saline was prepared and quantified at A_385 nm_ using an extinction coefficient of 16,760 M^-1^.cm^-1^. To measure the bis-ANS fluorescence emission upon binding to spike variants, we mixed 2.4 µL of bis-ANS (stock at 5 mg/mL) with a range of 15 to 60 µL depending on the spike variant to have a final concentration of 40 μg/mL bis-ANS and 20 μg/mL of spike protein in 300 µL reaction volume (Figure 3d). Bis-ANS fluorescence was recorded from 400 to 600 nm upon excitation at 360 nm.

Guanidine (Merck, #607-148-00-0) titrations were carried out in the presence of 20 μg/mL of each spike variant. Samples were prepared individually for each guanidine concentration (0, 0.01, 0.02, 0.03, 0.05, 0.06, 0.07, 0.08, 0.1, 0.15, 0.2, 0.25, 0.3, 0.4 0.5, 0.8, 0.9, 1, 1.3, 1.6, 1.9, 2.1, 2.5, 3, 3.5, 4, 4.5, and 5 M) and allowed to settle down at room temperature for 30 minutes before fluorescence acquisition. For these studies, the ancestral spike was either in PBS (pH 7.4) or 50 mM Tris-Cl containing 100 mM NaCl and 1 mM biotin at pH 8.0 (tris-based buffer). In this experiment, the ancestral protein, when tested in tris-based buffer, contained the TwinStrep® and the 8xHis purification tags (Figure 4b), while all other analyses (ancestral and variant spike proteins in the presence of PBS) were carried out with samples of tagless spike proteins, which had been treated with HRV-3C protease (Figures 4a and d-g).

Protein behavior as a function of pressure was investigated by applying pressure increments of 5,000 psi (∼344 bar) up to a maximum pressure of 3,500 bar. Upon each pressure increment, the pressure was allowed to stabilize for 5-min before bis-ANS fluorescence acquisition (Figure 6a). For these experiments, we mixed 5.5 µL of bis-ANS (stock at 5 mg/mL) with a range of 15 to 60 µL depending on the spike variant to have a final concentration of 25 μg/mL bis-ANS and 25 μg/mL of spike protein in 1,100 µL reaction volume. For light scattering (LS) measurements, samples were excited at 320 nm, and the emission was recorded from 300 to 340 nm (Figure 6b).

### Dynamic light scattering (DLS)

Measurements were performed on a Brookhaven 90Plus/BiMass multiple-angle particle sizing instrument. The autocorrelation curves C(τ) were fitted using multimodal size distribution (MSD), in which a non-negatively constrained least-square (NNLS) algorithm is used to produce an intensity or number-weighted distribution (Supplementary figure 5). Sample concentration was 50 μg/mL (final volume 100 μL) in PBS. Samples were allowed to rest for 20 minutes for thermal equilibration before acquisition at 25°C. C(τ) was acquired for 30 seconds for each run showing data as multiple replicates (Figure 3c).

### Thermodynamic parameters

*ΔG*_0M_^F-U^ values were obtained using the two-state equilibrium assumption of folding/unfolding (equations 2 and 3)

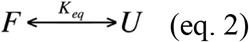

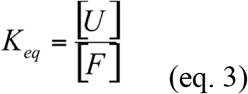

Considering [*U*] + [*F*] = 1, the fraction of unfolding (*α*) can be expressed as in equation 4 and correlated with *K*_*eq*_ as in equation 5

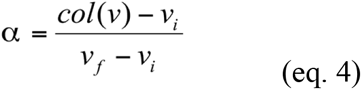

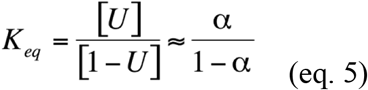

The fraction unfolded was calculated for each experimental condition, *col(v)* subtracted from its initial value (*v*_*i*_) and divided by the difference between the last (*v*_*f*_) and the first value. We calculated the free energy of denaturation extrapolated at 0 M GuHCl (Supplementary figure 8) as in equation 6

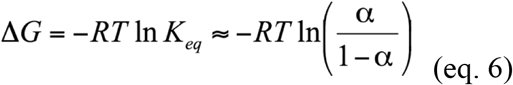

where ln[*α*/(1-*α*)] was obtained from the transition point’s intercept of the linear regression. R is the universal gas constant R (0.00198 kcal·K^-1^·mol^-1^), and T is temperature (298 K).

### Statistical Analysis

We used Minitab 17 software for statistical analysis. We performed a multiple comparison test for the half-maximal effective response (EC_50_), in which a one-way analysis of variance (ANOVA) was carried out, with a level of significance of 0.05. Model adequacy was examined by checking residual normality (Anderson-Darling test) and variance homogeneity (Bartlett test). The Post-Hoc Tukey test was implemented to determine which differences between pairs of EC_50_ means were significant.

## Acknowledgements

The authors thank Dr. B. Graham and Dr. K. Corbett (VRC/NIAID/NIH, USA) for sharing the pαH vector (BEI Resources #NR-52563) containing the gene construct for the ancestral spike protein. F.A.B.V. and L.R.C. gratefully acknowledge training on glycan analysis received from Dr. J. Cremata (Centro de Ingeniería Genética y Biotecnología, Cuba) (*in memoriam*). This work was supported by the Brazilian research funding agencies Empresa Brasileira de Pesquisa e Inovação Industrial (Embrapii), Conselho Nacional de Desenvolvimento Científico e Tecnológico (CNPq), Coordenação de Aperfeiçoamento de Pessoal de Nível Superior (CAPES) and Fundação de Amparo à Pesquisa do Estado do Rio de Janeiro (FAPERJ) to L.R.C.

The research was also supported by the Pew Charitable Trusts Foundation to G.A.P.d.O and the Carlos Chagas Filho Foundation for Research Support in the State of Rio de Janeiro (FAPERJ), grant 201.296/2021 to G.A.P.d.O and grants 210.008/2018, 202.840/2018, to J.L.S, the National Council for Scientific and Technological Development (CNPq), and the National Institute of Science and Technology for Structural Biology and Bioimaging (INCT), grants 465395/2014-7 and 402321/2016-2 to J.L.S.

## Authors contributions

H.R.S.A. performed fluorescence, CD, DLS and SEC experiments, data curation; T.M.L. purified spike variants; R.G.F.A. cultured, transfected and produced the HEK293 supernatants for spike purification; F.B.A.V. performed the HILIC-HPLC experiments; D.P.B.A. performed the ELISA experiments; F.F.M. aided on HEK293 culture; K.D.C. performed SDS-PAGE experiments; P.S-A, M.Q-V, J.S.G., and F.C.S.N. performed and analyzed mass spectrometry experiments; J.L.S., L.R.C., and G.A.P.d.O. conceptualization, data curation, funding acquisition, project administration, and íiting the manuscript; G.A.P.d.O. writing original draft, preparing figures, and supervising the research.

